# IAN: An Intelligent System for Omics Data Analysis and Discovery

**DOI:** 10.1101/2025.03.06.640921

**Authors:** Vijayaraj Nagarajan, Guangpu Shi, Reiko Horai, Cheng-Rong Yu, Jaanam Gopalakrishnan, Manoj Yadav, Michael H Liew, Calla Gentilucci, Rachel R Caspi

## Abstract

IAN is an R package that addresses the challenge of integrating, analyzing and interpreting high-throughput “omics” data, using a multi-agent artificial intelligence (AI) system. IAN leverages popular pathway and regulatory datasets (KEGG, WikiPathways, Reactome, GO, ChEA) and the STRING database for protein-protein interactions to perform standard enrichment analysis. The individual enrichment results are then used to generate insightful summaries, for each of the datasets, using a large language model (LLM) through a multi-agent architecture. These summaries are then contextually integrated and interpreted by the LLM, guided by carefully engineered prompts and grounding instructions, to provide insightful explanations, system overview, key regulators, novel observations etc. We demonstrate IAN’s potential to facilitate biological discovery from complex omics data, by reanalyzing two already published data and evaluating the results. We also show remarkable performance of IAN, in terms of avoiding hallucination. IAN package, along with installation instructions and example usage, is available on https://github.com/NIH-NEI/IAN.

## Introduction

The “Omics” scale technologies like genomics, transcriptomics, proteomics and metabolomics are helping researchers capture the whole of the biological system state at any given point of interest. The common workflow in most of these “Omics” studies, involve, generating system level molecular data between different conditions and then identifying molecules that are significantly differentially behaving between the conditions under study. This often results in a list of differentially behaving molecules (DEG - differentially expressed genes, for example in transcriptome studies). In a human system, this list usually contains at least a couple of thousands of genes, making it challenging to integrate it with other relevant data and study it as a system (Gomez-Cabrero et al., 2014; Lopez de Maturana et al., 2019). Since it is impossible to study and understand the relationship of each of those DEG’s with respect to the phenotype that is being studied, researchers often perform enrichment analysis to get a birds-eye-view idea on the system.

With numerous enrichment analysis tools and methods available, along with hundreds of gene sets to compare against, researchers have shown that none of the current methods are perfect and that most of them are biased and can produce skewed results (Nguyen et al., 2019). Performing the enrichment analysis using a multitude of resources, representing several biological system components like biochemical pathways, transcriptional factors, non-coding RNA targets, known disease-target associations, drug-target interactions etc., and then integrating them all together and trying to glean through the biological insights is not a trivial task, but a daunting one, if not an impossible feat. For example, the DAVID enrichment analysis platform (Sherman et al., 2022), produces results for more than 50 different datasets. Though the results from these 50 different datasets are integrated using a clustering approach, it is done mainly based on similarity between shared genes among the functional groups, without providing any intelligent insights into the system as a whole. Other popular tools like GSEA (Subramanian et al., 2005) (using hundreds of signatures from MSigDB) (Reimand et al., 2019), EnrichR (Chen et al., 2013) etc., also fall into the same realm as DAVID, with respect to their inability to comprehend and interpret all of the data, to provide any intelligent insights into the system. This is the problem we believe could be solved by LLMs.

In the past couple of years, LLMs have dramatically advanced the field of generative applications. In general, LLMs have significantly advanced the field of text generation, text summarization, translation, question answering, image generation, audio/video generation, code generation etc., (Tom B. Brown, 2020). Numerous LLM based applications are also being explored in the field of biomedical research and clinical settings, for their potential contributions toward drug design, drug discovery, sequence analysis, target discovery, clinical diagnosis, treatment recommendations, outcome predictions etc. (Ji et al., 2021; Lee et al., 2020; Thirunavukarasu et al., 2023). Unfortunately, the problem of hallucination is a major concern with LLMs use in the field of text generation (Farquhar et al., 2024). Researchers have proven that hallucination is an innate, inevitable limitation of LLMs, but are working towards identifying and using approaches, methods and tools that could be used to mitigate hallucination and improve LLMs capabilities (Kankanhalli, 2024). While researchers are working towards methods to address this concern in general, a recent study evaluated the use of LLMs for gene set enrichment and found that they were unsuitable as a replacement for standard enrichment analysis (Joachimiak et al., 2024). Other studies have also shown that LLMs are inferior in their ability to generate standards based scientific abstracts and scientific reports (Hwang et al., 2024; Wittmann, 2023). While LLMs have not yet impressed scientists with their general literature summarizing capabilities, providing them with a little help, by means of the retrieval augmentation, has proved to result in their state-of-the-art performance (Jin et al., 2024). Previous studies have also found that hallucinations are rare to non-existent when summarizing the gene sets through LLMs (Joachimiak et al., 2024).

Without tools to understand the system as a whole, researchers merely have been focusing on the top few hundred differentially expressed genes/transcripts, for example in transcriptomics studies, to propose disease mechanisms and to identify potential targets. Though tools like EnrichmentMap (Reimand et al., 2019) and Enrichr-KG (Evangelista et al., 2023) have enabled the possibility of looking at the enrichment results as a network, thereby understanding the relationship between enriched terms, they lack the ability to go beyond integrating results based on simple term similarity and known associations. The current tools lack the ability to “understand” and discover “insights” from the integrated data, like a human scientist would be able to do.

In this work, we have developed IAN, a multi-agent AI system, implemented as an opensource R package. Using publicly available data, we show how IAN is able to integrate, “understand” and discover “insights” from the omics data.

## Methods

### Architecture

IAN is implemented as an R package, integrating diverse data sources and analytical tools to facilitate systems-level biological discovery. IAN takes gene expression data (custom DEG lists, DESeq2 results and Seurat FindMarkers output) and performs enrichment analysis using clusterProfiler, ReactomePA, enrichR, and STRINGdb, generating a suite of pathway and network-based data/metrics. A multi-agent system then leverages these data/metrics, with each agent employing carefully crafted analysis instructions, as LLM prompts to summarize, categorize, and interpret specific aspects of the analysis. The resulting agent responses, combined with experimental design information, are used to construct a comprehensive augmented prompt, which is then employed for an integrated final LLM response. The integrated response is also used to generate an LLM derived system model. All the results and responses are parsed and presented, along with system insights, in a comprehensive HTML report. The IAN package was developed using R 4.4.1 on macOS Ventura platform. An overview of IAN’s system architecture is presented in Figure 1.

**Figure 1:**
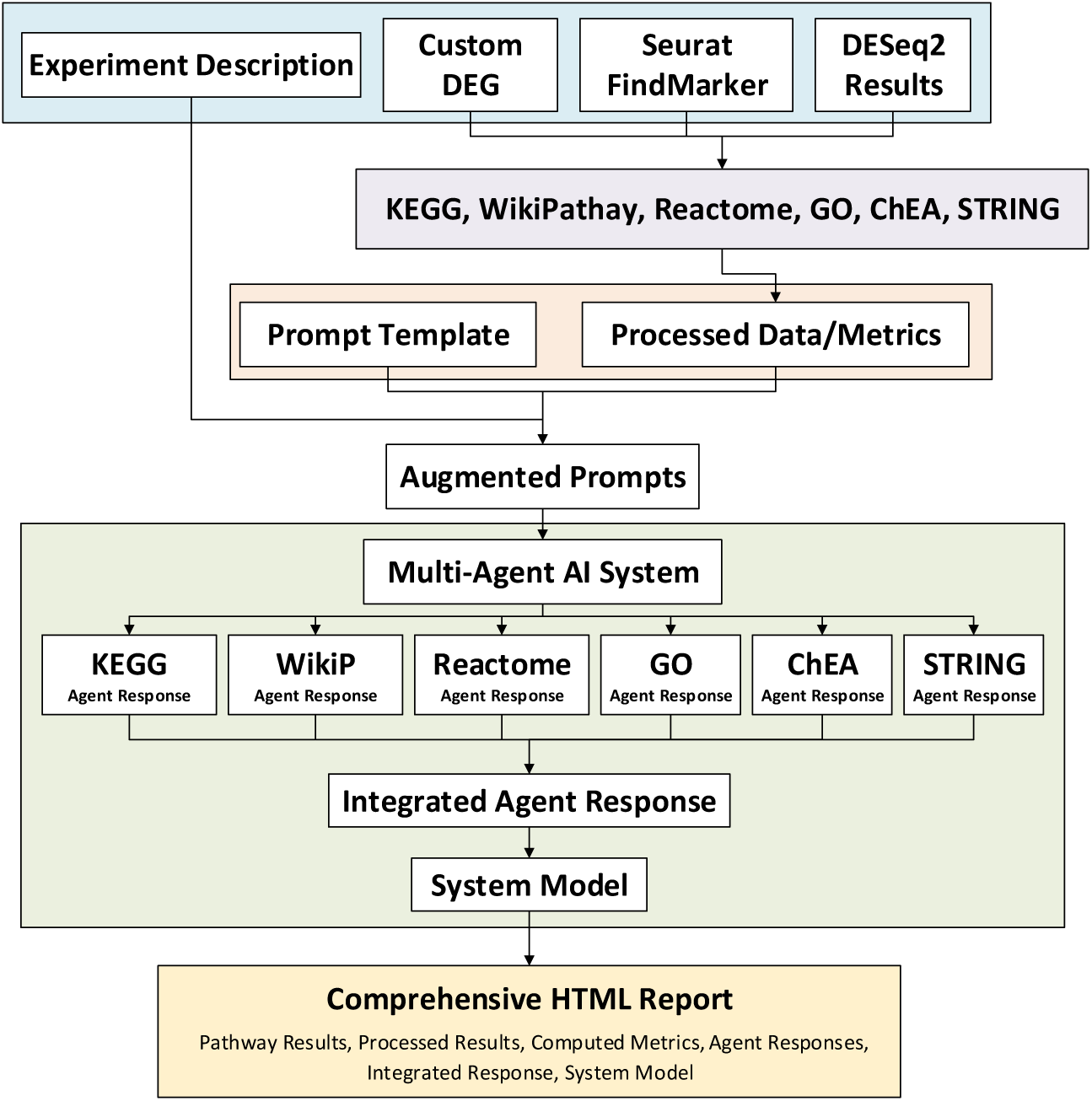
General system architecture of IAN

### Pathway Enrichment Analysis

To identify pathways involving the list of DEGs, IAN performs analysis using clusterProfiler’s (Yu et al., 2012) KEGG pathway enrichment, WikiPathways enrichment and Gene ontology enrichment, Reactome pathway enrichment (Yu & He, 2016) and enrichR’s ChEA transcription factor enrichment (Kuleshov et al., 2016). User provided DEGs are first mapped to ENTREZ IDs and Gene symbols, using clusterProfiler’s bitr function. The ENTREZ IDs are used as input for GO (Biological Process), KEGG, Reactome, and WikiPathways enrichment analysis. Gene symbols are used as input for ChEA. A p-value threshold of 0.05 is applied to filter the enrichment results for the top enriched terms. All enrichment results are further processed to map gene IDs to gene symbols and include only the enriched terms, enrichment scores and gene symbols for further processing. The processed enrichment results are also compared against each other to identify overlapping and unique genes, to provide additional input for the LLMs.

### Network Properties

For the given list of DEGs, IAN extracts all known interactions from the STRING database (Szklarczyk et al., 2023), with the default score as 0. The data is processed to remove duplicate interactions, convert string IDs to gene symbols and include only three columns (Protein1, Protein2, Combined Score). The extracted interactions are then sorted based on combined_score and the top 1000 interactions are kept for further analysis.

To identify key nodes within the protein-protein interaction network, IAN calculates several network properties for each gene, including degree (number of connections), betweenness (influence on network paths), closeness (proximity to other nodes), and centrality (connection to other influential nodes), using the igraph package (Nepusz, 2006). To integrate these diverse measures, each property is scaled to a mean of zero and standard deviation of one, ensuring equal contribution across different properties. A combined_properties_score is then calculated for each node by averaging its scaled property values, providing a holistic measure of network importance. Nodes are ranked based on the combined_properties_score, with higher scores indicating greater overall importance and influence within the system being studied.

### Multi-Agent System

The *R6* class system is used to implement IAN’s multi-agent core, creating the modular and reusable agent objects. Each agent (one for each of KEGG, WikiPathways, Reactome, GO, ChEA, STRING) is initialized with a specific LLM prompt and a corresponding prompt type, representing a distinct analytical task. The Environment class manages these six agents, orchestrating their execution using the *future* packages for parallel processing. The run_agents method distributes the prompts to the agents, retrieves responses from the Gemini API via a user-defined function, and handles retries and error conditions. The resulting agent responses are then aggregated and used to construct a final, combined prompt for a comprehensive system-level analysis.

### Large Language Model

IAN is optimized for Google’s “gemini-1.5-flash-latest” LLM, with a default temperature of 0 and the maximum token size of 8192. A five second delay is introduced with a maximum of three retries, to stay within the server overload limits. The user is expected to provide an API key to run IAN. We choose Google’s LLM server for their large context window availability.

### Report

Based on results and important concepts identified throughout the analysis, IAN generates a dynamic and comprehensive HTML report which is easy to share and user friendly to navigate. Tables of processed data, LLM summaries, plots and network graphs are included in the report, along with links to download all original results, processed results and full LLM reports.

### Implementation

Key functions included in IAN’s implementation is listed below. Detailed list of all the functions is provided in the package repository at https://github.com/NIH-NEI/IAN. Analysis instructions which form the core of the engineered prompts are also provided in the package repository. A combination of one-shot and chain-of-thoughts (Wei, 2022) approaches are used for prompt engineering.

- **IAN():** The central function orchestrating the entire “omics” analysis, integration and LLM-driven hypothesis generation workflow.
- **map_gene_ids():** Maps diverse gene identifiers to standardized ENTREZID and SYMBOL formats, ensuring compatibility across downstream analyses and LLM understanding.
- **perform_wp_enrichment()**: Identifies significantly enriched biological pathways using WikiPathways.
- **perform_kegg_enrichment()**: Identifies significantly enriched biological pathways using KEGG.
- **perform_reactome_enrichment()**: Identifies significantly enriched biological pathways using Reactome pathways.
- **perform_chea_enrichment()**: Identifies key transcription factors regulating the differentially expressed genes, revealing potential upstream regulators.
- **perform_go_enrichment()**: Identifies significantly enriched Gene Ontology (GO) terms.
- **perform_string_interactions()**: Extracts relevant protein-protein interactions from STRINGdb, and computes combined network properties scores for each of the differentially expressed genes.
- **create_combined_prompt()**: Integrates results from multiple enrichment analyses and network data into a single, comprehensive prompt for the LLM, enabling a holistic interpretation.
- **visualize_system_model()**: Generates an interactive network visualization of the system model, facilitating exploration of key relationships and potential mechanisms.

### Dependencies

IAN depends on Internet connectivity to run enrichment analysis and call the LLM. Critical R package dependencies include *dplyr* for data manipulation, *clusterProfiler*, *ReactomePA*, and *enrichR* for enrichment analyses, *STRINGdb* for protein-protein interaction data, and *igraph* and *visNetwork* for network analysis and visualization. Communication with the LLM is facilitated by *httr*, while parallel processing is enabled through *future*. Report generation leverages *rmarkdown*, and the overall architecture is built upon the *R6* class system. IAN can be installed and run on any operating system using R 4.4.1 and above.

### Evaluations

#### Dataset

We used two published RNA-Seq based transcriptomics datasets for evaluating IAN’s performance. The first study (Rosenbaum et al., 2021), provided a custom DEG list, generated by comparing uveitis patient transcriptomes against healthy controls. The second study (Zheng et al., 2022) provided raw RNA-Seq transcriptomics data generated by comparing Behcet’s disease (BD) patients and healthy controls. The BD RNA-Seq data was downloaded from NCBI GEO (<u>GSE198533</u>) and processed using the standard DESeq2 (Love et al., 2014) method.

#### Groundedness score

To assess the IAN’s ability to use the provided content in generating its responses, we calculated a groundedness score. We used this score as an indirect measure of hallucination by quantifying the extent to which the LLM’s output represents the provided input data. As the ground truth, we used the user provided list of differentially expressed genes and the significant terms identified from enrichment analysis. We then calculated the groundedness score (G) as the cardinality of the intersection between the ground truth (GT) and the genes/terms reported in the LLM’s response (LLM), divided by the cardinality of the genes/terms reported in the LLM’s response: G = |GT ∩ LLM| / |LLM|. This metric provides a quantitative assessment of the LLM’s ability to synthesize and integrate information from diverse sources, while still grounded in the provided content.

#### Contextual relevance

To evaluate the contextual relevance of the LLM’s responses in IAN, we measured the semantic similarity between the LLM-generated summaries and the input data containing enriched pathways and GO terms. We employed the transformer-based language model BERT (Bidirectional Encoder Representations from Transformers) to compare the LLM’s responses to the input data. Briefly, the text was tokenized, augmented with special tokens to delineate content boundaries, and chunked to accommodate BERT’s input length limitations. These chunks were then converted into input IDs, and relevant embeddings were extracted. The resulting chunked embeddings were combined and used to compute the cosine similarity score. This semantic similarity analysis was performed in a Python virtual environment using the transformers, torch, and sklearn.metrics.pairwise packages. The fully documented Python script used for this analysis is available on our project’s GitHub repository.

#### Human evaluation

To assess the quality and utility of the IAN-generated reports, we conducted a human evaluation comparing them against the standard manually curated analyses. We employed a Likert scale (1 to 5) to evaluate key aspects of the reports, including Accuracy, Relevance, Clarity, Trustworthiness, and Overall Satisfaction. To minimize ambiguity and ensure consistent scoring, we designed a scoring sheet with clear anchors and detailed descriptions for each score level. A panel of expert evaluators, comprising both senior scientists (four staff scientists) and junior researchers (four research fellows) from different laboratory sections assessed the reports based on the defined criteria. The template scoring sheet is provided as supplementary material.

#### Statistics

To quantify the expert human evaluation, we performed a Wilcoxon Signed-Rank test to assess whether the overall Likert scale scores were significantly skewed towards higher values, indicating positive user perception of the IAN-generated reports. Additionally, we employed the Wilcoxon Rank-Sum test to evaluate potential differences in Likert scale scores based on the level of expertise of the evaluators (Seniors Vs Juniors) and the specific dataset being analyzed (UV vs BD). Summary statistics, along with frequency distribution plots, were used to further characterize and understand the expert human evaluation scores.

## Results and Discussions

### R Package

IAN is available as an opensource R package at our project GitHub page: https://github.com/NIH-NEI/IAN. Installation instructions along with example usage, full results of the evaluated datasets (including all the AI generated responses, tools generated original results, prepared tools results, prepared metrics files, network files, experimental design information, parameters used, R session information) are provided at the project GitHub page. Detailed system analysis instructions and all function parameter descriptions are also shared through the GitHub page. Function level detailed documentation is provided through the package files.

After installing the necessary dependent R packages, IAN could be installed using the R command; `devtools::install_github("NIH-NEI/IAN")`

### Comprehensive Data Analysis

We analyzed two chronic autoimmune disease datasets (uveitis and Behcet’s disease) to understand IAN’s capabilities. We used results from both the analysis to evaluate IAN’s performance. Figure 2 shows a screenshot of the partial results report generated by IAN for the uveitis data analysis. All the individual input data and results files are also provided to the user through the Download section of the HTML results report (available from the project GitHub page).

**Figure 2:**
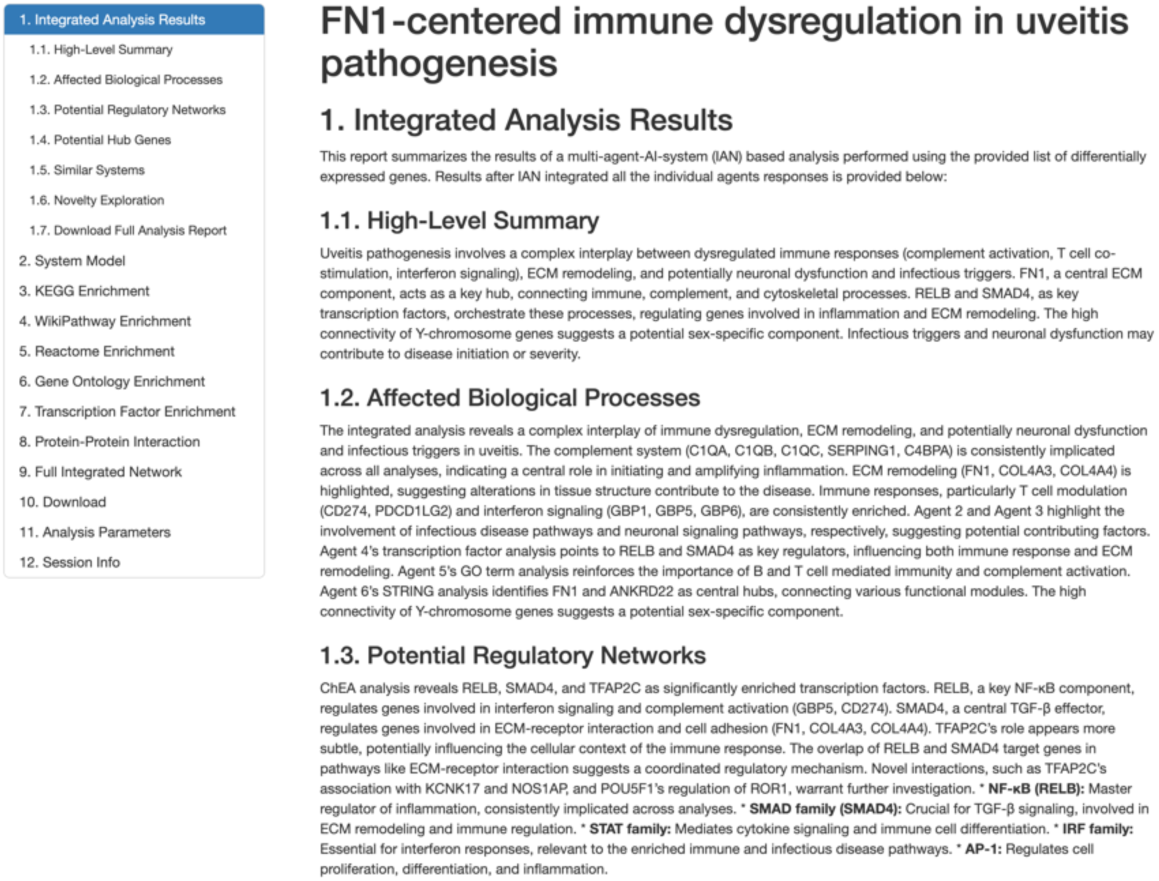
Screenshot showing the partial results as provided in IAN’s HTML report

#### RELB’s Influence on the Uveitis Phenotype

IAN’s HTML report starts with the LLM generated title, a high-level summary of the entire report, affected processes, potential regulatory networks, hub genes, similar systems, novelty exploration, system model overview, mechanistic model and hypothesis. These high-level results sections are followed by individual reports and summaries for each of the agents.

Here we briefly present and discuss the results from IAN’s analysis of uveitis dataset. Based on the comprehensive analysis performed by individual agents, IAN put forward this hypothesis – “*dysregulation of RELB, SMAD4, and ATF3 activity, potentially triggered by yet unidentified upstream factors, leads to altered expression of their target genes (GBP5, CD274, FN1), resulting in an amplified inflammatory response, impaired immune regulation, and altered tissue remodeling, contributing to the development and progression of uveitis*.”

The detailed analysis report shows that IAN performed a powerful analysis extracting and using more information and provided an in-depth comprehensive and actionable model from the same data that Rosenbaum used. The integrated summary confirms that IAN’s conclusions are strongly grounded in the input data and that it provides a highly plausible, meaningful, and useful model of uveitis pathogenesis. While Rosenbaum study (Rosenbaum et al., 2021) identified 10 genes that were also identified in Nussenblatt’s study (Li et al., 2008), to discuss their work, a comprehensive analysis and system level understanding of their own RNA-Seq data lacked, probably due to the inability and limitations of the tools that existed at the time of their study.

The LLM driven analysis of uveitis pathogenesis by IAN reveals a complex interplay of immune dysregulation, ECM remodeling, and potential neuronal/infectious involvement, with FN1 as a key hub. This model, built upon RNA-Seq data, identifies RELB, SMAD4 and ATF3 as key transcription factors orchestrating these processes. While IAN’s analysis identified several potential important mediators of uveitis pathogenesis, this discussion focuses specifically on the role of RELB, a key transcription factor implicated in immune regulation. Further investigation into the other hub genes and regulatory networks identified by IAN, such as FN1, CD274, SMAD4, ATF3 and the complement cascade, represents a promising avenue for future research to fully elucidate the complex mechanisms underlying uveitis.

IAN implicates RELB as a key regulator of genes involved in interferon signaling and complement activation, orchestrating the uveitis phenotype. Although there is no known literature that has reported any direct role for RELB in uveitis, there are plenty of studies that indirectly validates IANs inference, by supporting its general involvement in immune processes. For example, Tu et al.,(Tu et al., 2020) explore a novel TTP-RELB regulatory network for innate immunity gene expression, and Gupta et al.,(Gupta et al., 2019) show that RELB controls adaptive responses of astrocytes during sterile inflammation. A comprehensive review by Barnabei et al. (Barnabei et al., 2021) further strengthens these connections, highlighting RELB’s role in various immune cell types and its implications for autoimmunity and inflammation, noting that mutations in components of the NF-κB pathway, including RELB, have deepened our understanding of autoimmune disease. Furthermore, the presence of NFKBIB, IKBKG, and IKBKE genes in Nussenblatt’s (Li et al., 2008) list, provides indirect support for the involvement of NF-κB signaling pathways. Given its role in regulating inflammatory responses in other tissues, as highlighted by Elssner et al. (Elssner et al., 2019) showing increased RELB translocation in diseased human liver, it’s plausible that RELB contributes to the chronic inflammation and tissue damage observed in uveitis.

### Evaluation

#### Groundedness score

IAN scored a perfect 100% on the groundedness score, for a total of 7942 input tokens that we used to evaluate both the UV and BD datasets. The evaluated token categories included network property scores, all responses genes, final response genes, integrated network genes, system model genes, WikiPathway IDs, KEGG pathway IDs and Reactome pathway IDs. The High degree of groundedness demonstrates the reliability of IAN in generating insights that are directly supported by the provided input data. Detailed scores for different categories of both datasets evaluated are presented in the Supplemental Table 2.

#### Semantic similarity

Semantic similarity measured between the terms contained in all of the input contents and different responses from IAN showed a high degree of similarity across different comparisons, with an average score of 0.75. The strong similarity score demonstrates that IAN effectively captures and reflects the content of the input terms in its generated responses. Detailed semantic similarity scores for different categories, evaluating both the datasets, are presented in the Supplemental Table 3.

#### Expert human evaluation

The expert human evaluation of IAN’s analysis revealed strong performance across key metrics (Supplementary Table 4), with a mean accuracy of 4.75 (out of 5), high relevance (mean 4.50), excellent clarity (mean 4.63), and strong overall satisfaction (mean 4.75). While trustworthiness scored slightly lower (mean 4.25), these results collectively suggest that IAN’s AI-driven approach generates results that are not only accurate and relevant but also understandable and generally trusted by human experts. Furthermore, for all metrics evaluated (Accuracy, Relevance, Clarity, Trustworthiness and Overall Satisfaction), Wilcoxon rank-sum tests produced extremely high p-values, along with relatively small effect sizes, demonstrating that IAN’s results are reliably similar irrespective of who evaluated the reports and what dataset was used for evaluation. The effect sizes were all low when the evaluations were compared between the datasets. But, though the differences in the trustworthiness scores between the Senior and Junior groups were not significant, the effect size of 0.5 shows a potentially moderate difference between how much the Senior and Junior groups trust the AI generated reports. Given the paramount importance of trustworthiness in the field of AI and biomedical research, we opine that a large-scale study to further assess any disparities between senior and junior populations is warranted and that would be crucial to increase the adoption of the platform and paving way for discoveries by researchers of all age groups. Also, though there were ties in some scores, resulting in calculation of approximate p-values rather than exact p-values, we did not consider the ties to have any material effect on our conclusions, given the extremely high p-values and low effect sizes.

The Wilcoxon signed-rank test revealed a statistically significant skew towards higher scores (p-value = 0.0001264) indicating that the IANs report generally scored higher for both datasets, in both expert levels and across all metrics.

### Cost

During our development process, we made 1306 API calls to Google Gemini, spread across 30 days, costing us a pre-tax total of $3.48. We choose Google Gemini for the large context window as well as the low cost pay-as-you-go model.

## Limitations and future directions

IAN currently works only for Human and Mouse datasets, with ENSEMBL or ENTREZID or SYMBOLS, focusing on transcriptomics. We plan to include other model organisms and other ‘omics’ technologies like proteomics and metabolomics in our future version. Current version supports KEGG, WikiPathways, Reactome pathways, GO, ChEA and STRING data.

Future version would include other data sources and relevant computed metrics. While users can run IAN separately for up or down regulated genes, using the current version, future version would include options to specify and use the up/down information, along with the DEG. Current version of IAN is optimized for Google Gemini gemini-1.5-flash-latest model. Future versions would include other proprietary models, as well as opensource models like Llama3. We plan to implement several other features in the next version of IAN, including multi-dataset comparison, custom user ‘notes’ for the agents to focus on certain aspects and new analysis concepts in the summary report.

## Conclusions

In this work, we developed IAN, a multi-agent AI system to dig through ‘omics’ data, in an unprecedented way, combining user data with tools output and appropriate data metrics, to uncover deep insights into biological mechanisms, leveraging modern LLM’s superhuman-like reasoning capabilities. By analyzing and evaluating IAN’s performance on two published transcriptomics data, we demonstrate that IAN is capable of generating reliable results that are remarkably grounded in the user provided data and are conceptually similar to the content provided as input.

## Data availability

All of the input data and output data used in this work are provided in the project GitHub page (https://github.com/NIH-NEI/IAN/) for transparency and reproducibility.

Transcriptomics data evaluated by IAN are shared at Zenodo: https://doi.org/10.5281/zenodo.14974179.

## Acknowledgments

This work utilized the computational resources of the NIH HPC Biowulf cluster. The authors sincerely thank Drs. Richard Lee, Charles Egwuagu and Han-Yu Shih for their support and guidance throughout the related historical work. We also thank the NEI IRP program for funding this research.

## Author Contributions

VN conceptualized the study, developed the application, wrote the manuscript draft, performed evaluations and analyzed data. GS, RH, CY, JG, MY, MHL and CG performed evaluations and reviewed the manuscript. RRC contributed to the conceptualization, supervised the study, acquired funding, reviewed, edited and finalized the manuscript.

## Funding

This work was supported in part by the Intramural Research Program of the National Institutes of Health, National Eye Institute [EY000184, R01 EY032482].

## Supplementary Data

### Disclaimer: Use of Large Language Models

Authors declare that Google Gemini was used as an aid for literature review, to generate content ideas, to correct written text, as an algorithmic tool for research study, as an evaluation tool to identify anomalies in our code and text, as a coding assistant and for documentation drafting. The authors have reviewed all content generated by Gemini.

This supplementary file includes the following content:

**Supplementary Table 1:** Template scoring sheet (Likert scale) for Expert Human Evaluation.

| Score | Accuracy |
| --- | --- |
| 1 | The results and recommendations are completely inaccurate |
| 2 | The results and recommendations are slightly inaccurate |
| 3 | The results and recommendations are acceptable |
| 4 | The results and recommendations are mostly accurate |
| 5 | The results and recommendations are highly accurate |
|  | Relevance |
| 1 | The results and recommendations are not relevant to this study at all |
| 2 | The results and recommendations are slightly relevant to this study |
| 3 | The results and recommendations are relevant to this study |
| 4 | The results and recommendations are mostly relevant to this study |
| 5 | The results and recommendations are highly relevant to this study |
|  | Clarity |
| 1 | The results and recommendations are very difficult to understand |
| 2 | The results and recommendations are difficult to understand |
| 3 | The results and recommendations are clear |
| 4 | The results and recommendations are easy to understand |
| 5 | The results and recommendations are very easy to understand |
|  | Trustworthiness |
| 1 | Doesn't make sense, so doesn't seem trustable at all |
| 2 | Doesn't make much sense, so doesn't seem very trustable |
| 3 | Makes some sense, so seems somewhat trustable |
| 4 | Makes sense, so seems trustable |
| 5 | Makes perfect sense, so seems completely trustable |
|  | Overall Satisfaction |
| 1 | Completely useless, so would not recommend at all |
| 2 | Somewhat useless, would not recommend to most |
| 3 | Somewhat useful, would recommend with reservations |
| 4 | Useful, would recommend |
| 5 | Very useful, would highly recommend |

**Supplementary Table 2:** Ground truth categories and Groundedness score for UV and BD datasets analyzed by IAN.

|  | Input Tokens | **IAN-UV-Grounded** | Total reported | Groundedness score |
| --- | --- | --- | --- | --- |
| Network Properties Score | 96 | 9 | 9 | 100 |
| All Responses Genes | 316 | 62 | 62 | 100 |
| Final Response Genes | 316 | 22 | 22 | 100 |
| Integrated Network Genes | 316 | 15 | 15 | 100 |
| System Model Genes | 316 | 11 | 11 | 100 |
| WikiPathway IDs | 20 | 16 | 16 | 100 |
| KEGG Pathway IDs | 16 | 8 | 8 | 100 |
| **Average** |  |  |  | **100** |
| **Total** | **1396** |  |  |  |

|  | Input Tokens | **IAN-BD-Grounded** | IAN-Total reported | Groundedness score |
| --- | --- | --- | --- | --- |
| Network Properties Score | 453 | 13 | 13 | 100 |
| All Responses Genes | 1503 | 116 | 116 | 100 |
| Final Response Genes | 1503 | 21 | 21 | 100 |
| Integrated Network Genes | 1503 | 25 | 25 | 100 |
| System Model Genes | 1503 | 8 | 8 | 100 |
| WikiPathway IDs | 27 | 21 | 21 | 100 |
| Reactome Pathway IDs | 54 | 48 | 48 | 100 |
| **Average** |  |  |  | **100** |
| **Total** | **6546** |  |  |  |

**Supplementary Table 3:** BERT based semantic similarity scores for IAN generated reports.

|  | IAN-UV-Similarity |
| --- | --- |
| All Responses | 0.7688012 |
| Final Response | 0.65062845 |
| System Model | 0.71103364 |
| **Total/Average** | **0.71015443** |

|  | IAN-BD-Similarity |
| --- | --- |
| All Responses | 0.81039435 |
| Final Response | 0.7935405 |
| System Model | 0.77081585 |
| **Total/Average** | **0.791583567** |

**Supplementary Figure 1:**
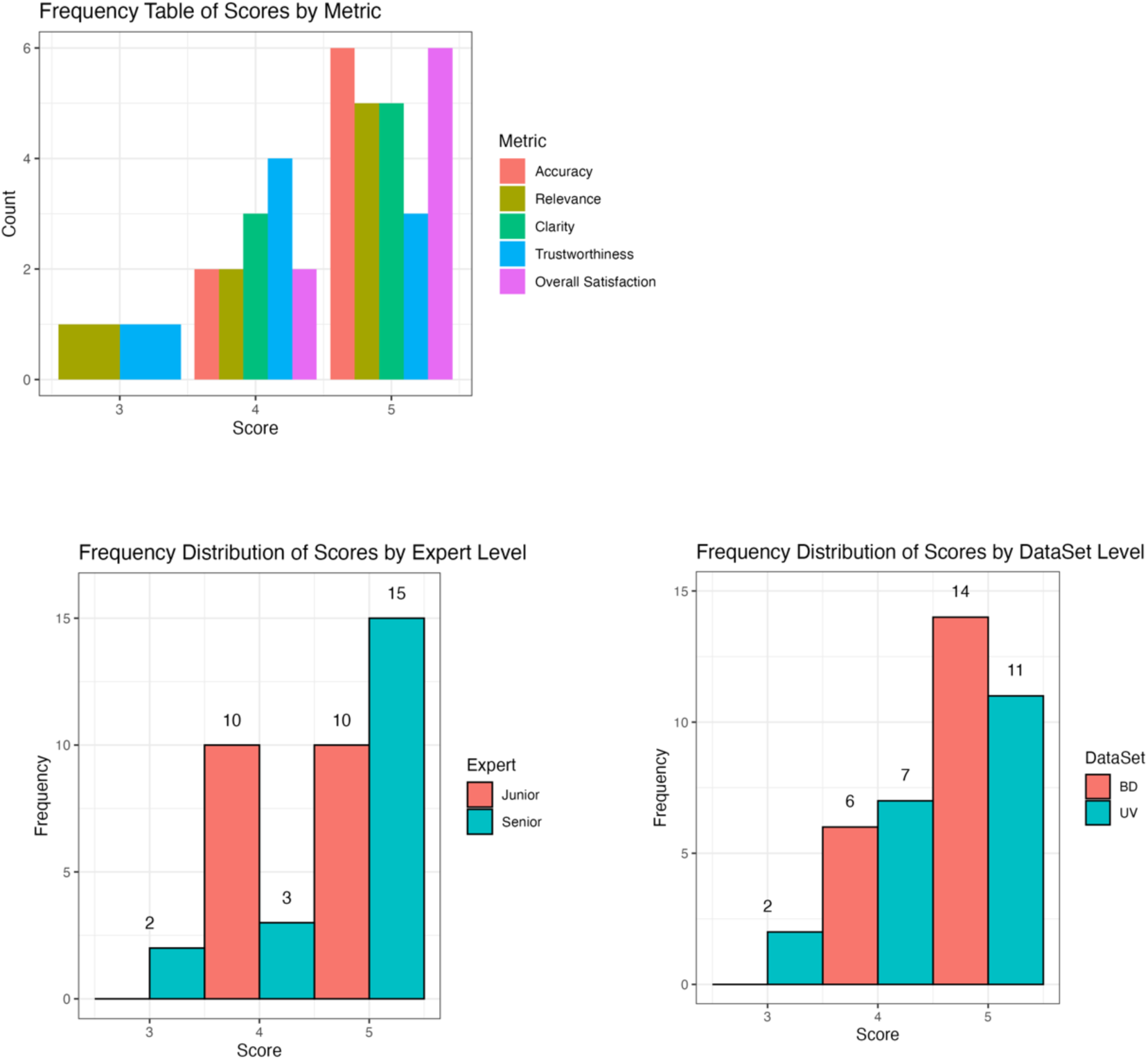
Frequency Tables and Frequency Distribution of expert human evaluation scores for IAN’s analysis reports.

**Supplementary Table 4:**
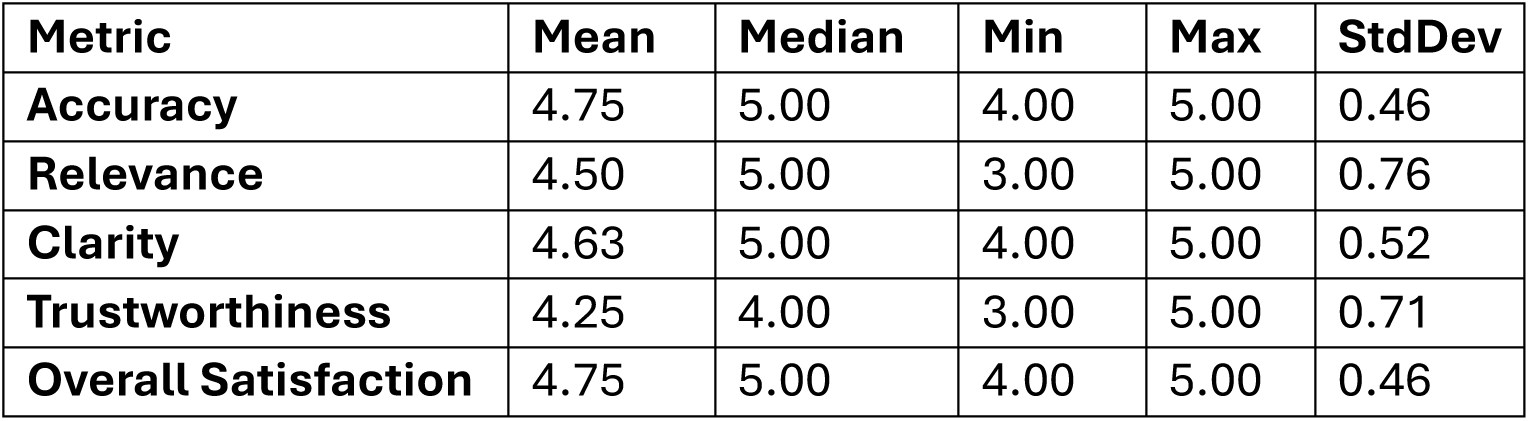
Summary statistics on human expert evaluation of IAN’s reports, using the Likert Scale of 1 to 5. 1 being worst performing, 3 average performing and 5 best performing.

**Supplementary Table 5:** The Wilcoxon Rank-Sum test showing no statistically significant differences in the IAN’s performance across datasets and expert levels.

| Datasets: UV vs BD |  |  |
| --- | --- | --- |
| Metric | P_value | Effect_Size |
| Accuracy | 1 | 0 |
| Relevance | 1 | -0.3125 |
| Clarity | 1 | -0.25 |
| Trustworthiness | 1 | -0.375 |
| Overall Satisfaction | 1 | 0 |

| Expert Level: Seniors vs Juniors |  |  |
| --- | --- | --- |
| Metric | P_value | Effect_Size |
| Accuracy | 1 | 0 |
| Relevance | 1 | 0.125 |
| Clarity | 1 | 0.25 |
| Trustworthiness | 1 | 0.5 |
| Overall Satisfaction | 1 | 0 |

## Notes

https://github.com/NIH-NEI/IAN

### Competing Interest Statement

The authors have declared no competing interest.

